# Comparative analysis of novel MGISEQ-2000 sequencing platform vs Illumina HiSeq 2500 for whole-genome sequencing

**DOI:** 10.1101/577080

**Authors:** Dmitriy Korostin, Nikolay Kulemin, Vladimir Naumov, Vera Belova, Dmitriy Kwon, Alexey Gorbachev

**Author notes:** Corresponding author (VB).

## Abstract

**Background:** MGISEQ-2000 developed by MGI Tech Co. Ltd. (a subsidiary of the BGI Group) is a new competitor of such next-generation sequencing platforms as NovaSeq and HiSeq (Illumina). Its sequencing principle is based on the DNB and cPAS technologies, which were also used in the previous version of the BGISEQ-500 device. However, the reagents for MGISEQ-2000 have been refined and the platform utilizes updated software. The cPAS method is an advanced technology based on cPAL previously created by Complete Genomics.

**Result:** In this paper, the authors compare the results of the whole-genome sequencing of a DNA sample from a Russian female donor performed on MGISEQ-2000 and Illumina HiSeq 2500 (both PE150). Two platforms were compared in terms of sequencing quality, number of errors and performance. Additionally, we performed variant calling using four different software packages: Samtools mpileaup, Strelka2, Sentieon, and GATK. The accuracy of single nucleotide polymorphism (SNP) detection was similar in the data generated by MGISEQ-2000 and HiSeq 2500, which was used as a reference. At the same time, a separate indel analysis of the overall error rate revealed similar FPR values and lower sensitivity.

**Conclusions:** it may be concluded with confidence that the data generated by the analyzed sequencing systems is characterized by comparable magnitudes of error and that MGISEQ-2000 can be used for a wide range of research tasks on par with HiSeq 2500.

## Background

The cPAL sequencing technology developed by Complete Genomics was first featured in a paper in 2009 [1]. In 2013, Complete Genomics was acquired by BGI (the Beijing Genomic Institute), and the technology has been subsequently refined [2]. In 2015, a new commercially available second-generation genome analyzer BGISEQ-500 was first announced [3]. Since then, the cPAL technology has undergone serious modifications.

The cPAS method was an important milestone in the evolution of this technology. The method utilizes fluorescently labeled terminated substrates. In the cPAS method, sequencing occurs as the DNA polymerase begins working with a primer (anchor) complementary to the single DNA strand [4]. DNA nanoballs (DNB) are 160,000 to 200,000-bp-long single-stranded DNA fragments made of replicated butt-joined copies of one of the original library DNA molecules, used for signal amplification. The copies are created in the process of rolling circle amplification of DNA rings, forming a library. Each DNB rests in a separate section of the patterned flow cell, which is ensured by its non-covalent binding to a charged substrate. The flow cell is a silicon wafer coated with silicon dioxide, titanium, hexamethyldisilazane, and a photoresistant material. DNBs are added to the flow cell and selectively binded to positively-charged aminosilanes in a highly-ordered pattern, allowing for the sequencing of a very high density of DNA nanoballs [1], [5].

The sequencing process itself consists of several steps, including the addition of a fluorescently labeled terminated nucleotide (sequencing by synthesis), the cleavage of a terminator during the synthesis process and the detection of the produced fluorescent signal [6], [7], [8]. We would like to emphasize that we were unable to find a detailed description of cPAS-based sequencing in the literature, nor were we able to figure out how it is implemented in MGISEQ-2000. However, a patent is available in the public domain that describes the application of the cPAS approach. In this patent, the sequencing process is described as using fluorescently labeled monoclonal antibodies that recognize unique chemical modifications of one of the four terminated dNTPs [9]. In any case, it is not currently possible to obtain full information on MGISEQ-2000 sequencing.

A paper was published two years ago, in which researchers used a reference genomic dataset obtained from GIAB to demonstrate that the BGISEQ-500 platform showed similar accuracy of SNP detection and slightly lower accuracy of indel detection compared to HiSeq 2500, [3]. Several recent studies have compared the performance of these two platforms in ancient DNA [10], metagenome [11] and microRNA [4] sequencing. In general, the quality of the data generated by BGISEQ-500 has proved to be satisfactory, although several of its characteristics were slightly worse than those of Illumina HiSeq 2500.

The Genome in a Bottle Consortium provides reference genomes for benchmarking [12]. By comparing the obtained genomic variants to a reference sequence, one can assess the accuracy/ sensitivity of the tested instrument and the corresponding bioinformatics pipeline for data analysis. In our study, we tested the suitability of the MGISEQ-2000 platform for the assessment of the mutational variability of embryonic cells. To do this, we used the genome of a Russian female egg donor and conducted a genome-wide analysis using two platforms: Illumina HiSeq 2500 and MGISEQ-2000. As HiSeq 2500 is a popular and well-described platform for genomic research, we decided to evaluate the overall error rate in order to understand whether we can use MGISEQ-2000 for the execution of our utilitarian tasks.

## Materials and methods

### Ethics approval and consent to participate

The research was carried out according to The Code of Ethics of the World Medical Association (Declaration of Helsinki). Written informed consent to participate and to publish these case details was obtained from the patient, and the study was approved by the Ethical Committee of Pirogov Russian National Research Medical University, Moscow, Russia.

### DNA preparation

A sample of genomic DNA was isolated from whole blood using phenol-chloroform extraction. Quality control was performed using agarose gel electrophoresis (degradation level) and the Qubit dsDNA BR Assay Kit (concentration measurement). The donor was a female resident of the Russian Federation.

### Preparation of a library for sequencing

#### MGISEQ-2000

The circularization procedure is essentially the denaturation and renaturation of the DNA library in the presence of excess amounts of a splint oligo (dephosphorylated at the 5’-end) that consists of inverted complementary sequences of adapters ligated to the library. In the process of renaturation with the splint oligo, an annular molecule is formed with a double-stranded structure in the adapter region containing a nick. The nick is sealed by a DNA ligase. Linear DNA library molecules are disposed of at the digestion stage using a mixture of nucleases that cleave linear molecules. A useful scheme was prepared by the MGI’s team [13].

The isothermal synthesis of nanoballs is carried out using the rolling circle amplification (RCA) mechanism and is initiated by the splint oligo. As a result, RCA forms a linear single-stranded DNA consisting of 300-500 repeats. A nanoball is a molecule compactly packed into a coil-like form 200-220 nm in diameter.

The procedure of loading of the nanoballs in the flow cell is simplified and automated: the flow cell has a patterned array structure that facilitates efficient loading (85.5% in our case), which does not depend on the accuracy of library dilution in the case of unordered cells (similar to, for example, Illumina MiSeq or HiSeq 2500). The nanoballs are loaded using a DNB Loader, a device similar to cBot (Illumina). The instrument and the reagents are prepared for sequencing in a way similar to that used for Illumina. Water and maintenance washes must be performed for MGISEQ-2000. The ready-to-use reagents are delivered in a cartridge that needs to be thawed prior to use. A flow cell for MGISEQ-2000 has four separate lanes and one surface, on which DNBs are immobilized.

We used MGIEasy Universal DNA Library Prep Set. 1000 ng of genomic DNA was fragmented using a Covaris ultrasonicator to achieve a length distribution of 100-700 bp with a peak at 350 bp. Size selection was performed using Ampure XP (Beckman). Library concentrations were measured using a Qubit; the amount of DNA used was 289 ng (procedure efficiency 29%). After that, an aliquot of 50 ng of the fragmentation product was transferred to a separate tube for end-repair and A-tailing. For ligation, the equimolarly mixed set of Barcode Adapters 501-508 was used. The ligation product was washed with Ampure XP, and seven PCR cycles were performed after that using primers complementary to the ligated adapters. After the washing of the library with Ampure XP, its concentration was measured using a Qubit. Before the annealing and circularization with splint oligo, the library was normalized to 330 ng in a volume of 60 μL. After linear DNA was digested, the concentration of ring DNA (0.997 ng/ μL) was measured using Qubit with the use of the ssDNA kit.

After RCA and formation of DNBs, the end product was measured using Qubit with the use of the ssDNA kit. The typical range of nanoball concentrations suitable for loading is 8-40 ng/ μL. In our case, the concentration was 20 ng/ μL. Nanoball loading was assisted by a DNB manual loader.

#### Illumina 2500

500 ng of genomic DNA was enzymatically fragmented by dsDNA Fragmentase (NEB). The library was prepared using the NEBNext Ultra II kit and indexes from the Dual Index Primers Set 2 (all New England Biolabs) according to the manufacturer’s instructions; amplification at the last sample preparation stage was performed in three PCR cycles.

MPS was carried out using the Illumina HiSeq 2500 in the Rapid Run mode (paired-end 150 bp dual indexing) with the use of the 500-cycle v2 reagent kit according to the manufacturer’s instructions.

#### Sequencing

Preparation of genomic libraries and sequencing using MGISEQ-2000 were carried out by our research group at the facilities of MGI Tech. in Shenzhen. Fastq files were generated as described previously using the zebracallV2 software provided by the manufacturer [3].

Library preparation and sequencing on HiSeq 2500 were carried out at the Center for Genome Technologies of Russian National Research Medical University. Fastq files were generated using the Basespace cloud software offered by the manufacturer (https://basespace.illumina.com/analyses/140691740/files/logs).

### Data analysis

The detailed description of the sequencing process and the protocols are provided in the S1 Additional file.

### Availability of data and material

Fastq files with WGS of E704 sample obtained using HiSeq 2500 and MGISEQ-2000 are available in SRA database (BioProject: PRJNA530191, direct link https://www.ncbi.nlm.nih.gov/Traces/study/?acc=PRJNA530191).

## Results

### Sequencing data summary

In this research, we analyzed two whole-genome datasets obtained by the sequencing of gDNA from a Russian female donor (hereinafter, we will call the sample E704). The donor’s genome was sequenced using two platforms: HiSeq 2500 by Illumina and new MGISEQ-2000 by BGI Complete Genomics that have similar performance characteristics. In the case of MGISEQ-2000, DNA was applied onto a separate lane of the flow cell. Sequencing was performed in a paired-end 150 bp mode. We recorded the amount of data generated by MGISEQ-2000 and calculated the average coverage. After that, we sequenced the donor’s genome using Illumina HiSeq 2500 in order to obtain a similar amount of data. General sequencing characteristics are presented in Table 1. The detailed description of library preparation is provided in the Materials and Methods section. We would like to note that we used different methods of DNA fragmentation for library preparation: fragmentation by ultrasound for E704-M and enzymatic fragmentation (dsDNA fragmentase) for E704-I. This fact is important for the interpretation of our results.

**Table 1.**
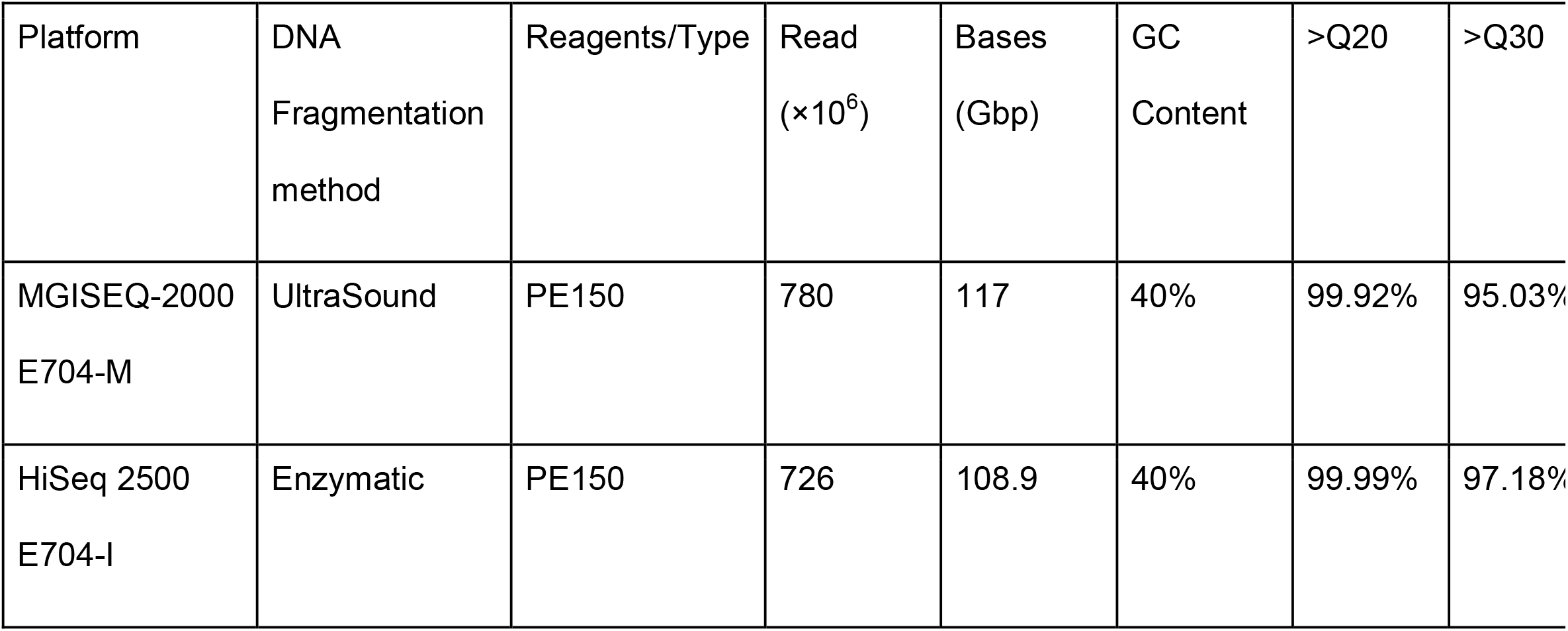
Summary of the dataset.

| Platform | DNA<br>Fragmentation<br>method | Reagents/Type | Read<br>(×10<sup>6</sup>) | Bases<br>(Gbp) | GC<br>Content | >Q20 | >Q30 |
| --- | --- | --- | --- | --- | --- | --- | --- |
| MGISEQ-2000<br>E704-M | UltraSound | PE150 | 780 | 117 | 40% | 99.92% | 95.03% |
| HiSeq 2500<br>E704-I | Enzymatic | PE150 | 726 | 108.9 | 40% | 99.99% | 97.18% |

As shown in Table 1, the size of the obtained dataset, as well as the characteristics of sequencing quality indicated that the datasets could be analyzed and compared. The comparison of the two datasets was unlikely to be skewed by the fact that different fragmentation methods were used [14].

### FastQC analysis

The next step in the comparison of the two datasets was to assess the quality of FastQ files using FastQC [15]. We also analyzed all individual FastQ files generated by paired-end sequencing (see *Materials and Methods*).

FastQC source file analysis demonstrated that the quality of data was acceptable and comparable for both platforms. K-mers were found at the start of the reads in the fastq files generated by MGISEQ-2000-based sequencing and at the end of the reads in the files generated by HiSeq 2500-based sequencing. A deviation from the normal GC-content was observed at the start of the reads in the HiSeq 2500 fastq files. Unremoved adapter sequences in both cases might explain the presence of K-mers. The abnormal GC-content could be a result of enzymatic fragmentation, which apparently causes a deviation from the random distribution pattern. Bearing that in mind, we decided to remove ten nucleotides from 5`-ends of each read in both MGISEQ-2000 and HiSeq 2500 fastq files. Further manipulations were carried out with 130-nucleotide-long fragmented reads. We also trimmed the adapter and other technical sequences (S1 Additional file), which allowed us to save more data and work with a higher average read length. This, however, was not crucial for our purposes, and we proceeded to the next steps of the comparative analysis. We merged all obtained fastq library files containing different barcodes so that each platform was represented by only a pair of fastq files with forward (R1) and reverse (R2) reads, respectively. After merging the fastq files, we repeated the quality assessment procedure using the FastQC service and found that the total data generated by both platforms was of acceptable quality and could be safely compared.

Figure 1 shows the assessment of quality of sequencing data by the FastQC service [15]. Data quality was acceptable for each of the nucleotide positions within a read for both MGISEQ-2000 and HiSeq 2500. However, the quality of data representing each position in the MGISEQ-2000 fastq file was slightly lower than in the HiSeq 2500 file and tended to gradually deteriorate towards the end of the read (although it was not lower than Q20). For HiSeq 2500-generated data, drops in quality below Q20 were observed only towards the very end of the read. For each nucleotide, the quality of MGISEQ-2000-based sequencing data gradually decreased after 50-60 cycles. In contrast, the total number of high-quality nucleotides was higher for HiSeq 2500 and remained on the same level until the last cycle. A similar picture can be seen in the graphs demonstrating the distribution of reads quality (Fig. 1c). The distribution was more uniform for Illumina, meaning that the average quality was higher. The quality of reads generated by the MGISEQ-2000-based sequencing was acceptable, as95% of all reads were above Q30. The GC-content was similar for both platforms (Fig. 1d); the distribution graphs are practically identical.

**Fig 1.**
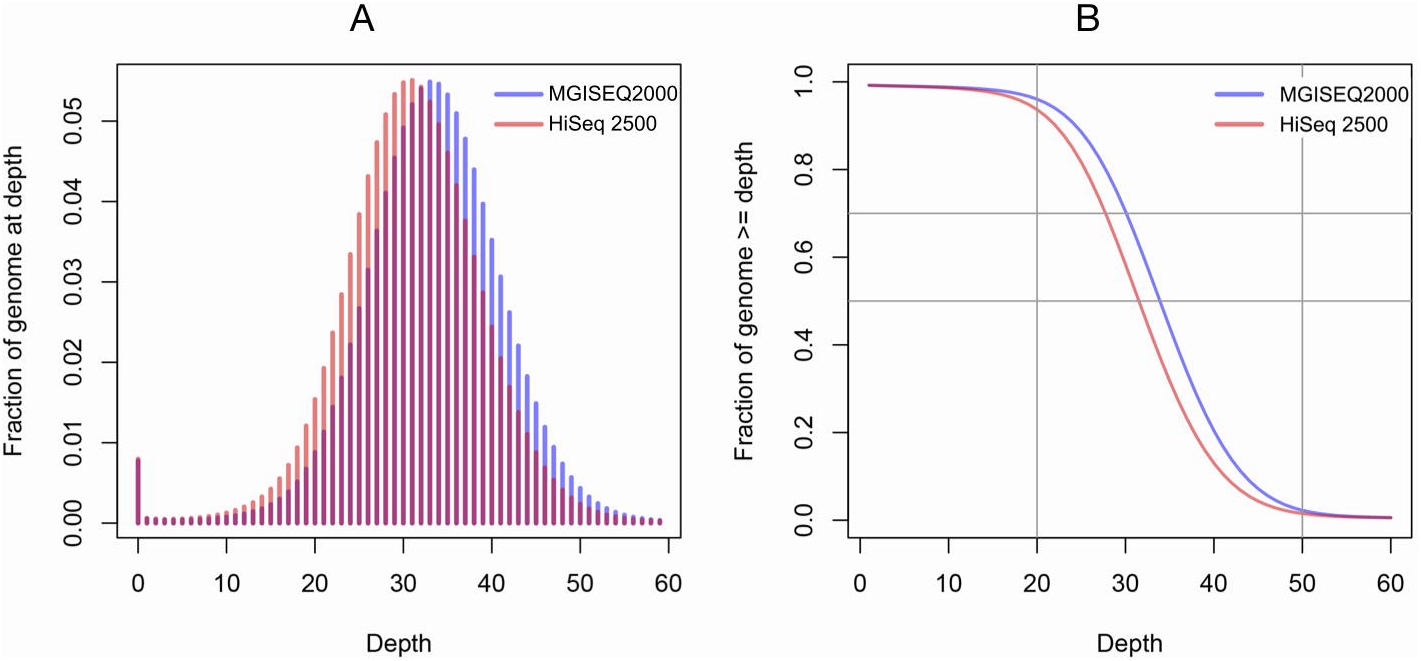
Post-filtering data quality control. (A), (B) Distribution of nucleotide quality parameters across reads. The presented data is for both MGISEQ-2000 (A) and HiSeq 2500 (B) platforms for forward (R1) and reverse (R2) reads, respectively. For each position in the reads, the quality scores of all reads were used to calculate the mean, median, and quantile values; therefore, the box plot can be shown. Overall quality score distribution for MGISEQ-2000 and HiSeq 2500 data (C). Distribution GC-content in the data generated by MGISEQ-2000 and HiSeq 2500 (D). FastQC [15] was used for the analysis.

### Reads mapping/ alignment and QC

The average coverage is an important characteristic of whole-genome sequencing, as are its distribution and variability. Figure 2 compares the average coverage distribution for MGISEQ-2000 and HiSeq 2500. The figure shows a slightly higher average coverage for MGISEQ-2000 (32.75X for MGISEQ-2000 versus 30.48X for HiSeq 2500). At the same time, the overall coverage distribution is highly uniform for both datasets (Inter-Quartile Range (IQR = 6)), suggesting good sequencing quality [18].

**Fig 2.**
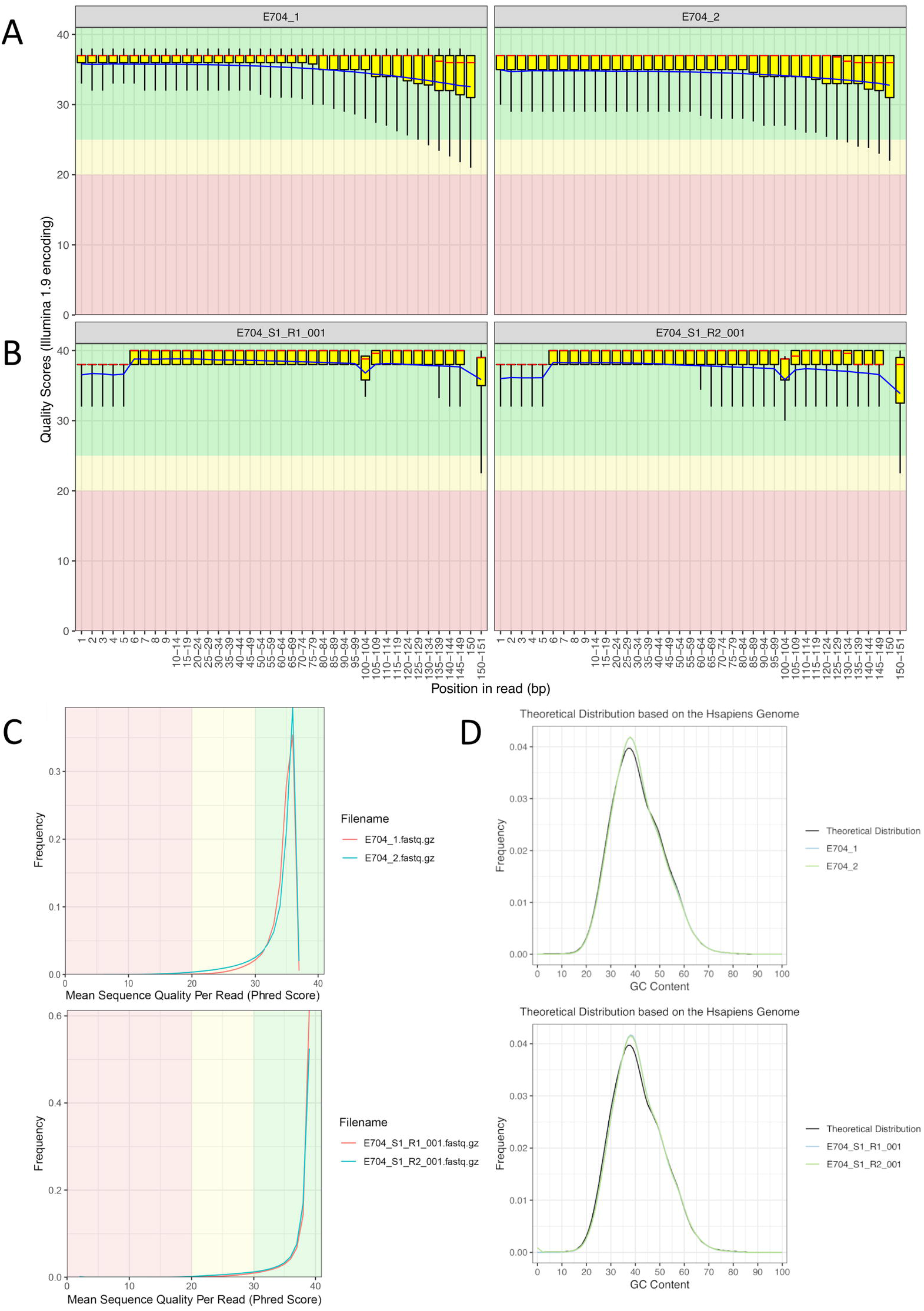
Analysis of the coverage distribution for MGISEQ-2000 and HiSeq 2500 with the use of the E704 sample. (A) A fraction of genome covered appropriate number of times. (B) A fraction of genome covered not less than the corresponding number of times. The analysis was performed using the R [16] and BEDtools [17] software packages.

The data presented in Figure 2 was obtained after the FastQC had been performed during the reads alignment. Therefore, the input data was similar in terms of the coverage distribution and the total reads number.

The filtered and trimmed reads were aligned to the reference genome, which was necessary to convert fastq files to BAM files. This was carried out using Burrows-Wheeler Aligner (BWA-MEM) with default settings recommended for the analysis of genomes sequenced on Illumina systems [19]. The quality of read alignment was assessed using the SAMtools software package and the bamstats software module [20, 21].

The quality of read alignment was acceptable for both platforms. The insert size for paired-end libraries corresponded to the theoretical size specified in the manufacturer’s protocol: 250 bp for Illumina HiSeq 2500 and 400 bp for MGISEQ-2000. The proportion of aligned reads was 99.9% for both BAM files.

Figure 3 presents the results of the analysis of read alignment to the reference genome. It is important that the frequency of random sequencing errors was much higher for MGISEQ-2000 and increased with the number of sequencing cycles.

**Fig 3.**
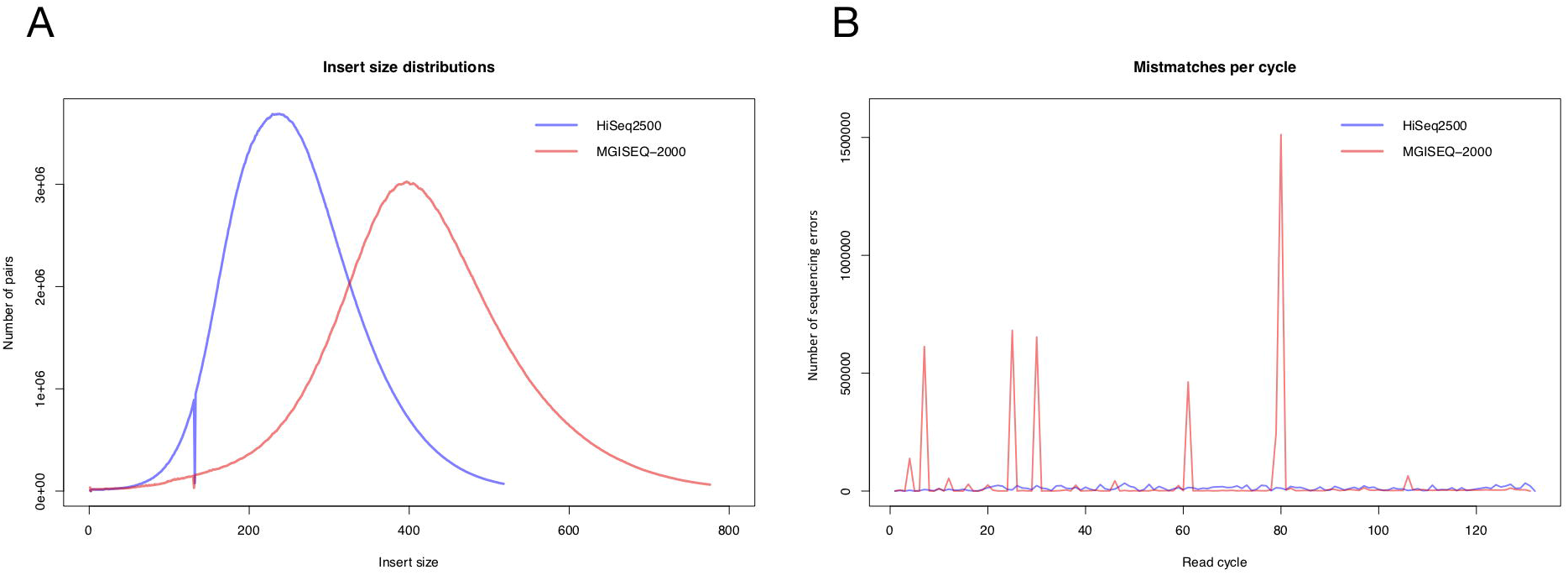
The results of the QC analysis of read alignment to the reference genome. (A) The distribution of insert length values between reads of the E704-I library (blue line) and the E704-M library (red line). (B) The number of random errors for HiSeq 2500 (blue line) and MGISEQ-2000 (red line). The alignment algorithm used is BWA-MEM [19]. QC analysis was performed using bamstats [20, 21].

### Variation calling and false positive/ negative ratio estimation

In order to further assess the quality of MGISEQ-2000 sequencing, as well as to understand the aspects of its potential use, the generated data was subjected to variant calling. After the data was aligned to the reference genome using BWA-MEM [19], the BAM file was modified using four different pipelines: Samtools [20, 21], Strelka2 [22], Sentieon [23], and GATK [24].

All software packages used to process the datasets generated by Illumina and MGI demonstrated similar performance in terms of computation speed, which is consistent with the results obtained for BGISEQ [25].

Alignment results are provided in Table 2; the table shows that both sequencing platforms performed similarly well. The duplication rate for E704-I was higher than for E704-M, amounting to 12.26%. This value, however, was calculated after we merged the fastq files with different barcodes and obtained from different lanes. In each individual fastq file, the duplication rate did not exceed 5-6% for both instruments (see S2 Additional file). Using Illumina HiSeq 2500, 16 separate fastq files (8 for + 8 rev) were generated. The number of fastq files obtained using MGISEQ-2000 was also 16, however, they represented a single flow cell, whereas Illumina’s files came from two different flow cells. Therefore, a higher duplication rate recorded for Illumina resulted from the use of two cells. Most likely, the probability of obtaining repeated reads from two independent flow cells is higher than from one cell. As the information in fastq files was summed up, it resulted in an additional 3-4% of duplicates for Illumina-generated data, compared to MGISEQ-2000.

**Table 2.** Mapping statistics for the datasets.

| Metrics | E704-M | E704-I |
| --- | --- | --- |
| Clean reads | 779784662 | 725927338 |
| Clean bases | 101372006060 | 94370553940 |
| HG19 length | 3095693983 | 3095693983 |
| Identified bases | 2921715981 | 2919239426 |
| Mapping rate | 99.85% | 99.93% |
| Unique rate | 90.83% | 87.20% |
| Duplication rate | 8.61% | 12.26% |
| Mismatch rate | 0.56% | 0.54% |
| Average Depth | 32.75 | 30.48 |
| Coverage at least<br>4x | 99.81% | 99.78% |
| Coverage at least<br>10x | 94.38% | 94.30% |
| Coverage at least<br>20x | 88.87% | 84.66% |

As it was not possible to conduct standard benchmarking procedures and determine error values in the reference genomic dataset during this study, we calculated error rates (False Positive, False Negative, etc.) in the E704-M dataset using E704-I as a reference. This approach cannot be used to assess the accuracy of the MGISEQ technology, however, it did allow us to conclude that the two compared technologies can be used interchangeably for similar tasks without significant loss of accuracy.

Figure 4 shows error rates determined with the use of different software packages. The best result was obtained by using Strelka2 [22]; below we will use the figures generated by this pipeline. Variant calling results are presented in the S2 Additional file. The magnitude of the total error (False Negative + False Positive) in the comparison of the samples E704-M and E704-I corresponded to the previously obtained results for BGISEQ500 and Illumina [https://blog.dnanexus.com/2018-07-02-comparison-of-bgiseq-500-to-illumina-novaseq-data/].

**Fig 4.**
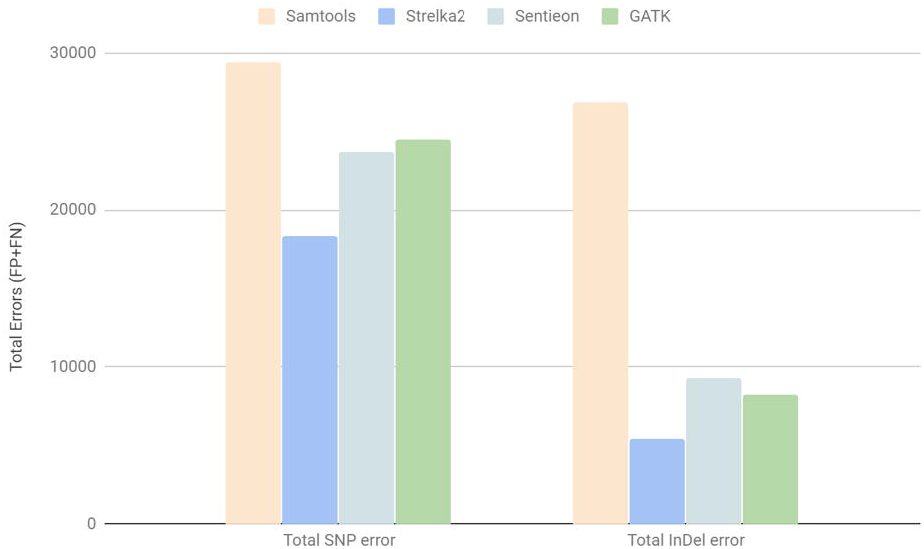
The total number of errors (the sum of FP and FN) for SNPs (total SNP error) and indels (total indel Error) detection that occurred in the course of genomic variants comparison of E704-M (A) and E704-I (B). Four software packages were used for variant calling: Samtool, Strelka2, Sentieon, and GATK. Baseline data is shown in the S2 additional file.

In total, over 3.7 million SNPs were detected in the datasets generated by each of the tested platforms. The E704-M sample contained 3,730,684 SNPs; the number of detected SNPs in the E704-I sample was comparable (3,719,768 SNPs). This data is shown in Table 3. In addition, we detected a similar Ti/ Tv ratio, which may indirectly indicate the sequencing accuracy.

**Table 3.**
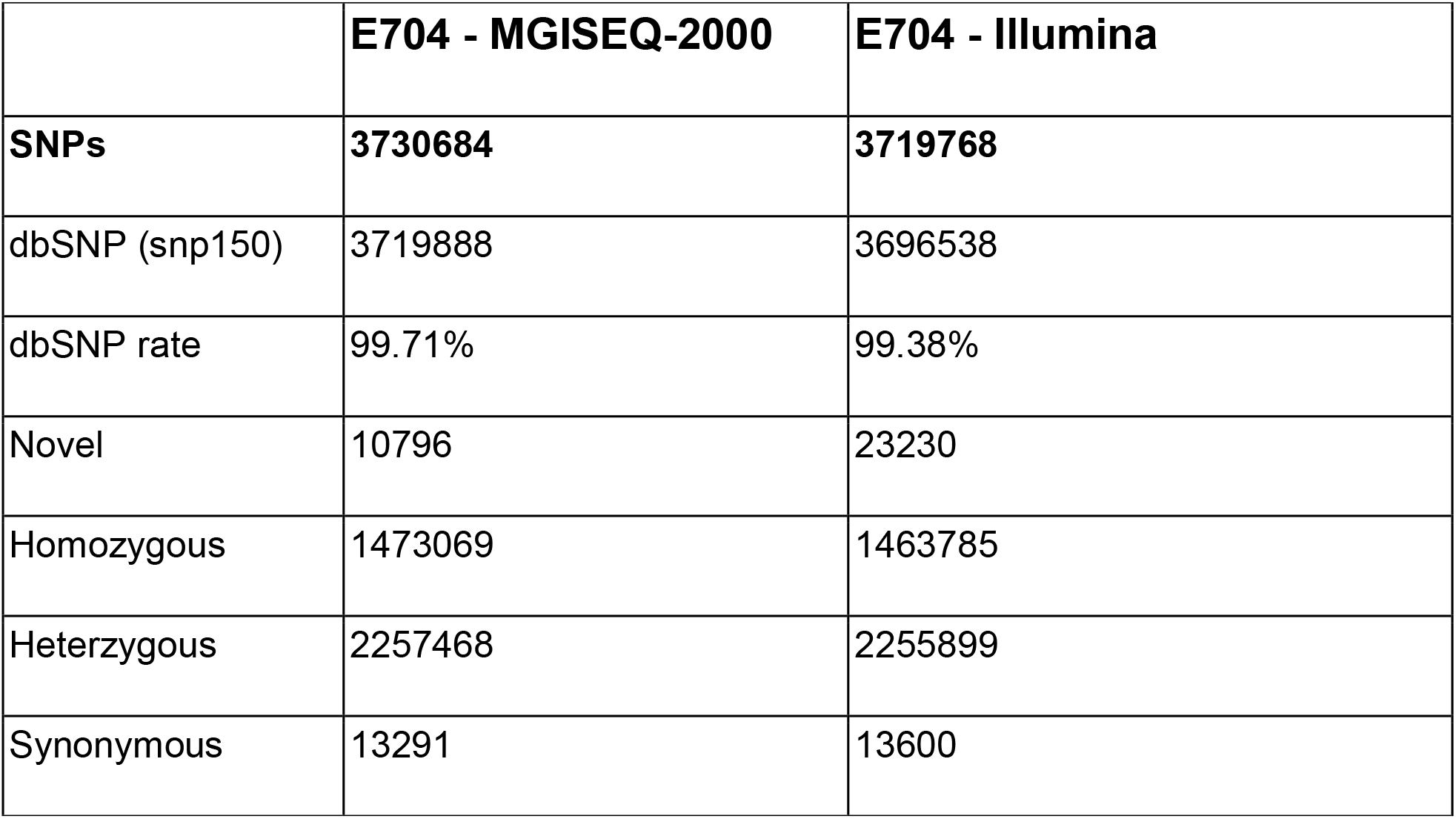
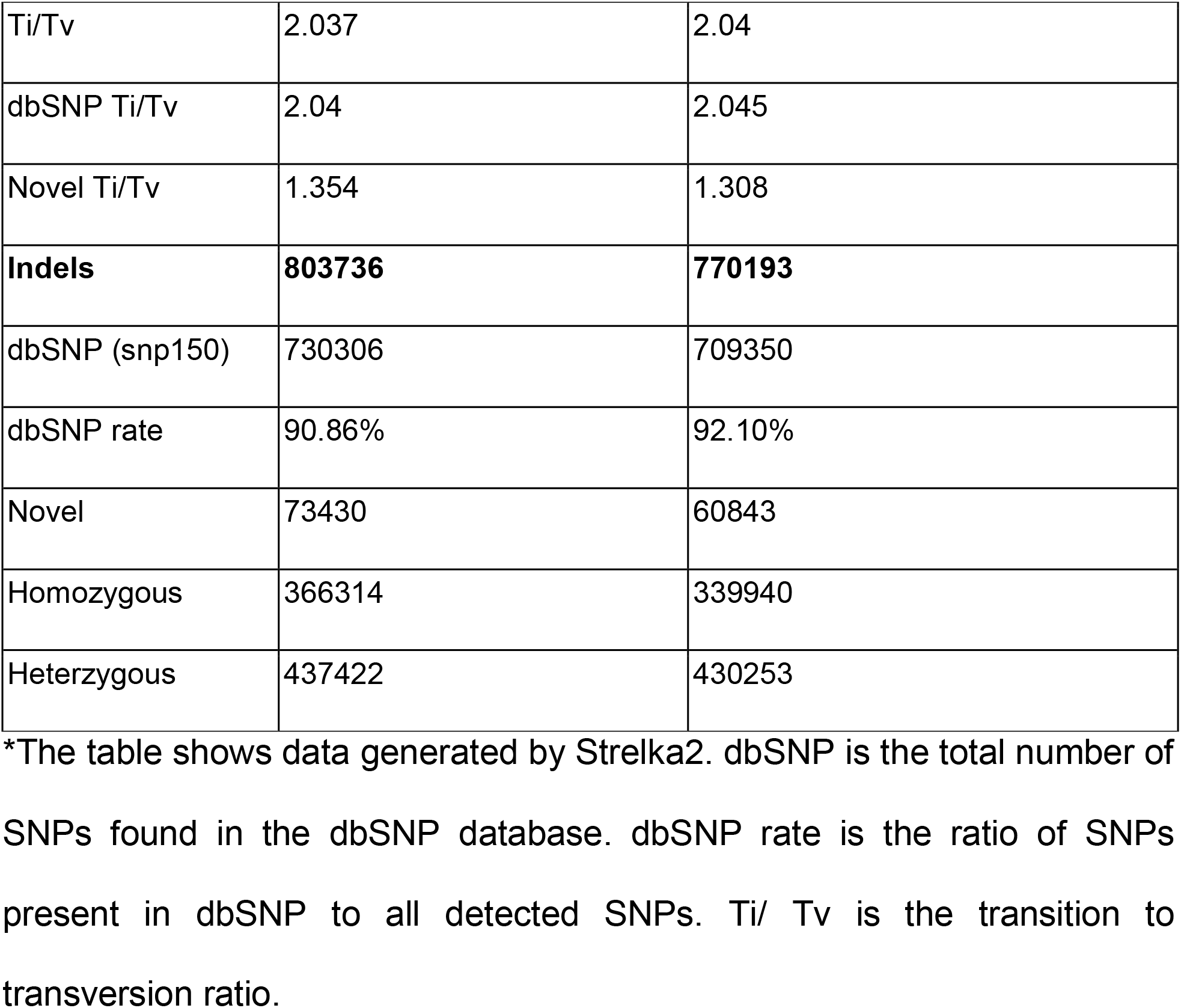
Variant calling statistics for the datasets*.

MGISEQ-2000 was able to detect slightly more indels (803,736) than HiSeq 2500 (770,193; see table 3). Generally, HiSeq 2500 performance was characterized by a slightly lower average coverage, which partly explains its indel detection rate. However, given that the dbSNP indel rate recorded by HiSeq 2500 was slightly higher (92.1% in E704-I versus 90.86% in E704-M), this may indicate a lower accuracy of indel detection by the MGISEQ-2000 platform. These observations are consistent with the previous findings for BGISEQ-500 [3].

To assess the accuracy of the detection of certain genomic variants, we chose the E704-I dataset as a reference for the E704-M sample. As a large number of such studies had been carried out for HiSeq 2500, we decided to determine the level of differences for a single genome. Sequencing using two different instruments allowed us to estimate their interchangeability/ similarity. We compared tested platforms using the HiSeq 2500 data as a reference, given that the permissible error rates for this technology have already been established by the Consortium. Further research using sequencing data from GIAB reference sample [12] to directly measure error rates for the detection of various mutations is needed.

We estimated the magnitude of various errors and calculated the F1-metric using vcf-compare (vcftools [26]) and snpeff [27]) for all detected SNPs.

Table 4 compares the variants obtained by variant calling using Strelka2; data generated by other software packages is presented in the S2 Additional file.

**Table 4.** Variant calling for E704-M versus E704-I.

|  |  | MGI vs Illumina |
| --- | --- | --- |
| **Identified bases (accessible genome)** |  | 2182021466 |
| **SNPs** | REF matches (full genome - VCF) | 2179423698 |
|  | All features in MGISEQ | 2597768 |
|  | REF matches (in VCF) | 2592230 |
|  | ALT matches (in VCF) | 2591850 |
|  | REF mismatches (in VCF) | 0 |
|  | ALT mismatches (in VCF) | 380 |
|  | In MGISEQ | 5538 |
|  | In reference | 12780 |
|  | In both | 2592230 |
|  | True Positive | 2592230 |
|  | False Positive | 5538 |
|  | True Negative | 2179423698 |
|  | False Negative | 12780 |
|  | TPR (Sensitivity, Recall) | 99.51% |
|  | TNR (Specificity) | 99.999746% |
|  | FNR | 0.49% |
|  | FPR | 0.000254% |
|  | PPV (Precision) | 99.79% |
|  | FOR | 0.00% |
|  | FDR | 0.21% |
|  | NPV | 100.00% |
|  | **F1-Metrics** | **99.65%** |
| **InDels** | REF matches for INDEL (VCF) | 2181793391 |
|  | All features in MGISEQ | 228212 |
|  | REF matches | 224595 |
|  | ALT matches | 223144 |
|  | REF mismatches | 842 |
|  | ALT mismatches | 1451 |
|  | In MGISEQ | 2775 |
|  | In reference | 2638 |
|  | In both | 225437 |
|  | True Positive | 225437 |
|  | False Positive | 2775 |
|  | True Negative | 2181793391 |
|  | False Negative | 2638 |
|  | TPR (Sensitivity) | 98.84% |
|  | TNR (Specificity) | 100.00% |
|  | FNR | 1.16% |
|  | FPR | 0.000127% |
|  | PPV (Precision) | 98.78% |
|  | FOR | 0.00% |
|  | FDR | 1.22% |
|  | NPV | 100.00% |
|  | **F1-Metrics** | **98.81%** |

As a result, using the “accessible genome” matrix, we discovered that the sensitivity of SNPs determination in the E704-M sample was 99.51% relative to the E704-I sample, with an FPR (false positive rate) value of 0.000254% (F1 metrics = 99.65%). For indels, the sensitivity was 98.84% (F1 metrics = 98.81%). It should be noted that although we did not perform a comparison with the reference sequence, the level of convergence of genotypes for MGISEQ-2000 and Illumina Hiseq2500 was high enough for both the accessible genome and the complete sequence of the read genome. This demonstrated that the MGISEQ-2000 sequencing had higher accuracy compared to previously obtained data for BGISEQ-500 [3]. This data is shown in Table 4.

## Discussion

We compared two genomic datasets generated by Illumina HiSeq 2500 and MGISEQ-2000-based sequencing. As part of our study, we aimed to understand whether MGISEQ-2000 could be used for the whole-genome sequencing of embryos, SNP detection and other tasks that our laboratory performs.

Our study demonstrated that MGISEQ-2000 provided datasets possessing characteristics similar to the data generated by the “gold standard” of the NGS analysis — the Illumina platform. Given a comparable amount of output data (101.37Gb for MGISEQ and 94.37Gb for Illumina), the average coverage for the two sets was comparable: 32.75X for MGISEQ-2000 versus 30.48X for HiSeq250; the coverage distribution patterns were almost identical (Figure 1).

The analysis demonstrated that the studied instruments provide similar sequencing quality. The existing differences can be explained by the specifics of the preliminary steps of library preparation and are not the result of the features of the sequencing techniques themselves.

Four different pipelines were used to perform variant calling. The detection rate of genomic variants in the two datasets was similar. The computational time required to process the obtained data was comparable for all software packages and all datasets used. The performance of Strelka2 was characterized by the lowest number of errors (Figure 4).

The quality of data obtained with MGISEQ-2000 was inferior in several respects to that generated by Illumina HiSeq 2500. Specifically, the frequency of random sequencing errors, the percentage of quality reads, and the accuracy of indel detection were higher for HiSeq 2500. However, the magnitude of those differences is small and insignificant for most research tasks. Last but not least, sequencing costs are an important factor for the laboratories. To our knowledge, the MGISEQ-2000 platform is comparable to NovaSeq in terms of costs, however, it requires a smaller number of samples per run.

## Conclusions

The newly-developed sequencer MGISEQ-2000 from BGI Group can be used as a fully-featured alternative to Illumina sequencers in whole-genome surveys (variant calling, indels detection). Raw data quality had equal metrics. Differences between two platforms that we found in the processes of variant calling and indel detection were negligible.

## List of abbreviations

bp: base-pair
cPAS: combinatorial Probe-Anchor Synthesis
dATP: deoxyadenosine triphosphate
dTTP: deoxythymidine triphosphate
DNBs: DNA nanoballs
FNR: false negative rate
FPR: false positive rate
FN: false negative
FP: false positive
GIAB: Genome in A Bottle
MPS: Massive Parallel Sequencing
PCR: polymerase chain reaction
PE150: pair-end 150 bp
SNPs: Single Nucleotide Polymorphisms
indels: insertions and deletions
WGS: Whole Genome Sequencing
WBC: White Blood Cell
DK: Dmitriy Korostin
VB: Vera Belova
DKw: Dmitry Kwon
NK: Nikolay Kulemin
AG: Alexey Gorbachev
VN: Vladimir Naumov

## Authors’ contributions

DK had designed the project. DKw and DK conducted sample preparation and sequencing library construction. VB, DKw and DK conducted sequencing. NK, VN and AG conducted data analysis. DK and AG wrote the manuscript. All authors have read and approved the manuscript.

DK - Dmitriy Korostin

VB - Vera Belova

DKw - Dmitry Kwon

NK - Nikolay Kulemin

AG - Alexey Gorbachev

VN - Vladimir Naumov

## Acknowledgements

We thank the Center for Precision Genome Editing and Genetic Technologies for Biomedicine (Moscow) for the genetic research methods.

## References

1. Drmanac R, Sparks AB, Callow MJ, Halpern AL, Burns NL, Kermani BG, et al. Human genome sequencing using unchained base reads on self-assembling DNA nanoarrays. Science. 2010;327:78–81.

2. Specter M. The Gene Factory [Internet]. The New Yorker. The New Yorker; 2013 [cited 2018 Dec 25]. Available from: https://www.newyorker.com/magazine/2014/01/06/the-gene-factory

3. Huang J, Liang X, Xuan Y, Geng C, Li Y, Lu H, et al. A reference human genome dataset of the BGISEQ-500 sequencer. Gigascience. 2017;6:1–9.

4. Fehlmann T, Reinheimer S, Geng C, Su X, Drmanac S, Alexeev A, et al. cPAS-based sequencing on the BGISEQ-500 to explore small non-coding RNAs. Clin Epigenetics. BioMed Central; 2016;8:123.

5. Chrisey LA, Lee GU, O’Ferrall CE. Covalent attachment of synthetic DNA to self-assembled monolayer films. Nucleic Acids Res. 1996;24:3031–9.

6. Canard B, Sarfati RS. DNA polymerase fluorescent substrates with reversible 3’-tags. Gene. 1994;148:1–6.

7. Mitra RD, Shendure J, Olejnik J, Edyta-Krzymanska-Olejnik, Church GM. Fluorescent in situ sequencing on polymerase colonies. Anal Biochem. 2003;320:55–65.

8. Tsien RY, Ross P, Fahnestock M, Johnston AJ. Dna sequencing [Internet]. Patent. 1991 [cited 2018 Dec 25]. Available from: https://patents.google.com/patent/CA2044616A1/en

9. Drmanac R, Drmanac S, Li H, Xu X, Callow MJ, Eckhardt L, et al. Stepwise sequencing by non-labeled reversible terminators or natural nucleotides [Internet]. US Patent. 2018 [cited 2018 Dec 25]. Available from: https://patentimages.storage.googleapis.com/46/2d/9b/5a6013e915f9b7/US20180223358A1.pdf

10. Mak SST, Gopalakrishnan S, Carøe C, Geng C, Liu S, Sinding M-HS, et al. Comparative performance of the BGISEQ-500 vs Illumina HiSeq 2500 sequencing platforms for palaeogenomic sequencing. Gigascience. 2017;6:1–13.

11. Fang C, Zhong H, Lin Y, Chen B, Han M, Ren H, et al. Assessment of the cPAS-based BGISEQ-500 platform for metagenomic sequencing. Gigascience. 2018;7:1–8.

12. Zook JM, Chapman B, Wang J, Mittelman D, Hofmann O, Hide W, et al. Integrating human sequence data sets provides a resource of benchmark SNP and indel genotype calls. Nat Biotechnol. 2014;32:246–51.

13. Oligos and primers for BGISEQ&MGISEQ NGS system [Internet]. [cited 2019 April 03]. Avaliable from: http://en.mgitech.cn/include/upload/kind/file/20181108/20181108161128_5692.pdf

14. Knierim E, Lucke B, Schwarz JM, Schuelke M, Seelow D. Systematic comparison of three methods for fragmentation of long-range PCR products for next generation sequencing. PLoS One. 2011;6:e28240.

15. Babraham Bioinformatics - FastQC A Quality Control tool for High Throughput Sequence Data [Internet]. [cited 2019 Jan 28]. Available from: http://www.bioinformatics.babraham.ac.uk/projects/fastqc/

16. Ripley BD. The R project in statistical computing. MSOR Connections The newsletter of the LTSN Maths, Stats & OR Network. 2001;1:23–5.

17. Quinlan AR, Hall IM. BEDTools: a flexible suite of utilities for comparing genomic features. Bioinformatics. 2010;26:841–2.

18. Sequencing Coverage for NGS Experiments [Internet]. [cited 2019 Jan 28]. Available from: https://www.illumina.com/science/education/sequencing-coverage.html

19. Li H, Durbin R. Fast and accurate short read alignment with Burrows-Wheeler transform. Bioinformatics. 2009;25:1754–60.

20. Li H. A statistical framework for SNP calling, mutation discovery, association mapping and population genetical parameter estimation from sequencing data. Bioinformatics. 2011;27:2987–93.

21. Li H, Handsaker B, Wysoker A, Fennell T, Ruan J, Homer N, et al. The Sequence Alignment/Map format and SAMtools. Bioinformatics. 2009;25:2078–9.

22. Kim S, Scheffler K, Halpern AL, Bekritsky MA, Noh E, Källberg M, et al. Strelka2: fast and accurate calling of germline and somatic variants. Nat Methods. 2018;15:591–4.

23. Weber JA, Aldana R, Gallagher BD, Edwards JS. Sentieon DNA pipeline for variant detection - Software-only solution, over 20× faster than GATK 3.3 with identical results [Internet]. PeerJ PrePrints; 2016 Jan. Report No.: e1672v2. Available from: https://peerj.com/preprints/1672/

24. McKenna A, Hanna M, Banks E, Sivachenko A, Cibulskis K, Kernytsky A, et al. The Genome Analysis Toolkit: a MapReduce framework for analyzing next-generation DNA sequencing data. Genome Res. 2010;20:1297–303.

25. Carroll A. Comparison of BGISEQ 500 to Illumina NovaSeq Data [Internet]. Inside DNAnexus. 2018 [cited 2019 Feb 15]. Available from: https://blog.dnanexus.com/2018-07-02-comparison-of-bgiseq-500-to-illumina-novaseq-data/

26. Danecek P, Auton A, Abecasis G, Albers CA, Banks E, DePristo MA, et al. The variant call format and VCFtools. Bioinformatics. 2011;27:2156–8.

27. Cingolani P, Platts A, Wang LL, Coon M, Nguyen T, Wang L, et al. A program for annotating and predicting the effects of single nucleotide polymorphisms, SnpEff: SNPs in the genome of Drosophila melanogaster strain w1118; iso-2; iso-3. Fly. 2012;6:80–92.

