## Additional file 1 for "Comparative analysis of novel MGISEQ-2000 sequencing platform vs Illumina HiSeq 2500 for whole-genome sequencing"

**S1 Additional file. Data analysis protocol**

Raw data was provided in FASTQ format and verified by **FastQC**.

For MGISEQ, a significant amount of k-mers were found in the 5’ area of reads (10 nucleotides).

For Illumina, a significant deviation from the GC composition was also found at the first 10 nucleotides.

It was decided to cut 10 nucleotides at the beginning of 5’end for both sequences.

They were cut off by the **cutadapt -u 10 -u -10** command

Specific **trimmomati**c trimming was performed.

**java -jar /data/tools/progs/Trimmomatic-0.36/trimmomatic-0.36.jar PE -threads 2 -phred33 -trimlog cut_special/E704_ILL/E704_ILL_cut.log F.fq.gz R.fq.gz Paired_F.fq.gz Unpaired_F.fq.gz Paired_R.fq.gz Unpaired_R.fq.gz ILLUMINACLIP:<list of adapters.txt>:2:30:10 LEADING:3 TRAILING:3 SLIDINGWINDOW:4:15 MINLEN:50 &**

FASTQ after cut off were passed through **FastQC** again, all test were passed correctly.

**BWA MEM** alignment was performed without additional parameters.

**bwa mem -t 2 hg19_karyo_order.fa forward_reads.fastq.gz reverse_reads.fastq.gz | awk -F'\t' '{if($4 != 0){print;}}' | samtools view -Shu - | samtools sort -o result.sorted.bam
samtools index result.sorted.bam**

Alignment quality were tested using **samtools stats**.

**samtools stats -r hg19_karyo_order.fa result.sorted.bam > stats_list.txt**

**plot-bamstats -p bamstats/ stats_list.txt**

The amount of aligned reads were counted by reverse analysis method from bam file.

**samtools fastq E704.sorted.bam | wc -l**

The total amount of lines should be divided by 4 (FASTQ definition).

Coverage was monitored by the **bedtools genomecov** package

**bedtools genomecov -bga -i result.sorted.bam > result.genomicstats.bed**

Then the total statistics is calculated by the following script (PHP7.2 CLI):

*<?php*

*if ($argc != 3) die ("count_cov_stats.php *.fa *.bed");*

*$faifile = file("{$argv["1"]}.fai");*

*$bedfile = fopen("{$argv["2"]}", "r");*

*$qual_arr = array("0" => 0, 4, 10, 20, 30, 40, 50, 70, 100, 200, 500, 1000, 10000);*

*while ($line = fgets($bedfile, 100000)) {*

*if (substr($line, 0 ,1) == '#') continue;*

*$ll = explode("\t", trim($line));*

*$gg = explode("-", trim($ll["3"]));*

*if (!isset($count["{$ll["0"]}"]["{$gg["0"]}"])) $count["{$ll["0"]}"]["{$gg["0"]}"] = 0;*

*$count["{$ll["0"]}"]["{$gg["0"]}"] = $count["{$ll["0"]}"]["{$gg["0"]}"] + $ll["2"] - $ll["1"];*

*}*

*echo ("Chrom\tFull_length");*

*for ($t = 1; isset($qual_arr["$t"]); $t++) echo ("\tPerc {$qual_arr["$t"]}x");*

*echo ("\n");*

*$sum_all = 0;*

*for ($gg = 0; isset($faifile["$gg"]); $gg++) {*

*$kl = explode("\t", $faifile["$gg"]);*

*$sum_all = $sum_all + $kl["1"];*

*$sum_cov_chr = 0;*

*echo ("{$kl["0"]}\t{$kl["1"]}");*

*for ($t = 1; isset($qual_arr["$t"]); $t++) {*

*if (!isset($sum_cov["{$qual_arr["$t"]}"])) $sum_cov["{$qual_arr["$t"]}"] = 0;*

*if (!isset($count["{$kl["0"]}"]["{$qual_arr["$t"]}"])) $count["{$kl["0"]}"]["{$qual_arr["$t"]}"] = 0;*

*$sum_cov_chr = $sum_cov_chr + $count["{$kl["0"]}"]["{$qual_arr["$t"]}"];*

*}*

*for ($t = 1; isset($qual_arr["$t"]); $t++) {*

*$perc = round($sum_cov_chr*100/$kl["1"], 2);*

*echo ("\t{$perc}%");*

*$sum_cov["{$qual_arr["$t"]}"] = $sum_cov["{$qual_arr["$t"]}"] + $sum_cov_chr;*

*$sum_cov_chr = $sum_cov_chr - $count["{$kl["0"]}"]["{$qual_arr["$t"]}"];*

*}*

*echo ("\n");*

*}*

*echo ("Full Genome\t$sum_all");*

*for ($t = 1; isset($qual_arr["$t"]); $t++) {*

*$perc = round($sum_cov["{$qual_arr["$t"]}"]*100/$sum_all, 2);*

*echo ("\t{$perc}%");*

*}*

*echo ("\n");*

*?>*

SNP and InDel calling was made by four different tools:

Using an uncalibrated BAM file:

1) **samtools mpileup + bcftools -g** (in gvcf) with a minimum total coverage of 10 (the rest is the default), the quality control was made by vcftools.

Calling was produced splitted into chromosomes to speed the process up.

**samtools mpileup -u -t DP -r chr1 -f hg19_karyo_order.fa result.sorted.bam | bcftools call -m -g 10 | vcf-annotate -f +/d=10/-a | grep -E "^#|PASS" --line-buffered > result_chr1.gvcf &**

Files were merged with **cat** tool.

2) **strelkav2**

**python configureStrelkaGermlineWorkflow.py --bam result.sorted.bam --ref hg19_karyo_order.fa --runDir *run_dir***

**python *run_dir*/runWorkflow.py**

Using a recalibrated file:

3) **Sentieon** (own recalibration in the protocol)

*REFERENCE="hg19_karyo_order.fa"*

*SENTIEON="path_to_sention"*

*export SENTIEON_LICENSE="our_license"*

*($SENTIEON bwa mem -M -R '@RG\tID:GROUP_NAME\tSM:SAMPLE_NAME\tPL:PLATFORM' -t 2 $REFERENCE ${1} ${2} || echo -n 'error' ) | $SENTIEON util sort -r $REFERENCE -o ${3}.sorted.bam -t 2 --sam2bam -i -*

*$SENTIEON driver -t 2 -r $REFERENCE -i ${3}.sorted.bam --algo GCBias --summary ${3}_GC_SUMMARY.txt ${3}_GC_METRIC.txt --algo MeanQualityByCycle ${3}_MQ_METRIC.txt --algo QualDistribution ${3}_QD_METRIC.txt --algo InsertSizeMetricAlgo ${3}_IS_METRIC.txt --algo AlignmentStat ${3}_ALN_METRIC.txt*

*$SENTIEON plot GCBias -o ${3}_GC_METRIC.pdf ${3}_GC_METRIC.txt*

*$SENTIEON plot MeanQualityByCycle -o ${3}_MQ_METRIC.pdf ${3}_MQ_METRIC.txt*

*$SENTIEON plot QualDistribution -o ${3}_QD_METRIC.pdf ${3}_QD_METRIC.txt*

*$SENTIEON plot InsertSizeMetricAlgo -o ${3}_IS_METRIC.pdf ${3}_IS_METRIC.txt*

*$SENTIEON driver -t 2 -i ${3}.sorted.bam --algo LocusCollector --fun score_info ${3}_SCORE.txt*

*$SENTIEON driver -t 2 -i ${3}.sorted.bam --algo Dedup --rmdup --score_info ${3}_SCORE.txt --metrics ${3}_DEDUP_METRIC.txt ${3}_DEDUP.bam*

*$SENTIEON driver -t 2 -r $REFERENCE -i ${3}_DEDUP.bam --algo Realigner ${3}_REALIGNED.bam*

*$SENTIEON driver -t 2 -r $REFERENCE -i ${3}_REALIGNED.bam --algo QualCal ${3}_RECAL_DATA.TABLE*

*$SENTIEON driver -t 2 -r $REFERENCE -i ${3}_REALIGNED.bam -q ${3}_RECAL_DATA.TABLE --algo QualCal ${3}_RECAL_DATA.TABLE.POST*

*$SENTIEON driver -t 2 --algo QualCal --plot --before ${3}_RECAL_DATA.TABLE --after ${3}_RECAL_DATA.TABLE.POST ${3}_RECAL_RESULT.CSV*

*$SENTIEON plot QualCal -o ${3}_BQSR.pdf ${3}_RECAL_RESULT.CSV*

*$SENTIEON driver -t 2 -r $REFERENCE -i ${3}_REALIGNED.bam -q ${3}_RECAL_DATA.TABLE --algo Haplotyper --emit_mode gvcf ${3}_VARIANT.vcf*

*$SENTIEON driver -r $REFERENCE --algo GVCFtyper -v ${3}_VARIANT.vcf ${3}_GVAR.vcf*

4) **GATK** (own recalibration in the protocol)

*REFERENCE="hg19_karyo_order.fa"*

*GATK="GenomeAnalysisTK-3.2-2/GenomeAnalysisTK.jar"*

*MILLS="GenomeAnalysisTK-3.2-2/Mills_and_1000G_gold_standard.indels.hg19.sites.vcf"*

*PHASE="GenomeAnalysisTK-3.2-2/1000G_phase1.indels.hg19.sites.vcf"*

*PICARD="picard/picard.jar"*

*#picardtools*

*java -jar $PICARD AddOrReplaceReadGroups I=*нерекалиброванный BAM* O=grupped.bam RGID=4 RGLB=lib1 RGPL=illumina RGPU=unit1 RGSM=20*

*samtools index grupped.bam*

*java -jar $PICARD MarkDuplicates I=grupped.bam O=rmdup_grupped.bam M=dup_metrics.txt REMOVE_DUPLICATES=false*

*#BAMindex*

*samtools index rmdup_grupped.bam*

*#Realignment*

*java -jar $GATK -T RealignerTargetCreator -R $REFERENCE -I rmdup_grupped.bam -o intervals.bed -known $MILLS -known $PHASE*

*java -jar $GATK -T IndelRealigner -R $REFERENCE -I rmdup_grupped.bam -targetIntervals intervals.bed -o realigned.bam -known $MILLS -known $PHASE*

*#BaseRecalibrator(BQSR)*

*samtools index realigned.bam*

*java -jar $GATK -T BaseRecalibrator -nct 2 -R $REFERENCE -I realigned.bam -o recalibration.bed -knownSites $MILLS -knownSites $PHASE*

*java -jar $GATK -T PrintReads -R $REFERENCE -I realigned.bam -BQSR recalibration.bed -baqGOP 30 -o recalibrated.bam*

*#Reduce bam*

*samtools index recalibrated.bam*

*java -jar $GATK -T ReduceReads -R $REFERENCE -I recalibrated.bam -o final.bam*

*samtools index final.bam*

*#Genotype*

*java -jar $GATK -T HaplotypeCaller -R $REFERENCE -I recalibrated.bam -stand_call_conf 30.0 -stand_emit_conf 10.0 -o main.vcf*

5) **samtools mpileup + bcftools -g** (in gvcf) started from the **GATK** recalibrated.bam file with a minimum total coverage of 10 (the rest is the default), the quality control was made by vcftools.

Calling was produced splitted into chromosomes to speed the process up.

**samtools mpileup -u -t DP -r chr1 -f hg19_karyo_order.fa recalibrated.bam | bcftools call -m -g 10 | vcf-annotate -f +/d=10/-a | grep -E "^#|PASS" --line-buffered > result_chr1.gvcf &**
Files were merged with **cat** tool.

Only samtools pipeline provided the GVCF file (all other tools - only VCF). So we used our own script to transform GVCF in VCF (PHP7.2 CLI)

*<?php
$file = fopen($argv["1"], "r");
while ($line = fgets($file, 4096)) {
if (substr($line,0,1) == '#') echo $line;
else {
$llg = explode("END=", $line);
if (isset($llg["1"])) continue;
$ll = explode("\t", $line);
if ($ll["4"] == '.') continue;
echo $line;
}
}
fclose($file);
?>*

All VCF files were split into SNP and InDel parts using

**grep (-v) INDEL variants.vcf**

All statistics were count by **vcf-compare** and **snpshift**.

**vcf-compare vcf1.vcf.gz vcf2.vcf.gz**

**java -Xmx1g -jar SnpSift.jar concordance -v vcf1.vcf vcf2.vcf**

As an additional comparison, for some callers, VCF-files were reduced using 1000genomes accessible regions and ClinVar matrices.

**ftp://ftp.1000genomes.ebi.ac.uk/vol1/ftp/release/20130502/supporting/accessible_genome_masks/20141020.strict_mask.whole_genome.bed**

**ftp://**[**ftp.ncbi.nlm.nih.gov/pub/clinvar/vcf_GRCh37/clinvar_20190225.vcf.gz**](http://ftp.ncbi.nlm.nih.gov/pub/clinvar/vcf_GRCh37/clinvar_20190225.vcf.gz)

The **tabix -R *bed-file* *vcf-file before filtering* > *vcf-file after filtering*** was used to filter files from step 10.

The specified files were also compared by **vcf-compare** and **snpshift**.

**vcf-compare vcf1.vcf.gz vcf2.vcf.gz**

**java -Xmx1g -jar SnpSift.jar concordance -v vcf1.vcf vcf2.vcf**

As an addition - the R code to build figure 1 graphics.

*#### /bdt_cover_hist_all.txt*

*col1 = rgb(0,0,1,alpha=0.5)*

*col2 = rgb(1,0,0,alpha=0.5)*

*bdt_cover_hist_all <- read.delim("bdt_cover_hist_all.txt", header=FALSE, stringsAsFactors=FALSE)*

*gcov = bdt_cover_hist_all*

*maxd = 400*

*plot(gcov[1:maxd,2], gcov[1:maxd,5],type='h', col=col1, lwd=3,xlab="Depth", ylab="Fraction of regions at depth")*

*plot(gcov[1:maxd,2], gcov[1:maxd,5],type='h', col=col2, lwd=3,xlab="Depth", ylab="Fraction of regions at depth",xlim=c(0,40))*

*gcov_cumul = 1 - cumsum(gcov[,5])*

*plot(gcov[2:(maxd+1),2], gcov_cumul[1:maxd],col=col1, type='l', lwd=3,xlab="Depth", ylab="Fraction of regions >= depth",ylim=c(0,1.0))*

*abline(v = c(0,50,100,200,300), col = "gray60")*

*abline(h = c(0,1,0.5), col = "gray60")*

*plot(gcov[2:(maxd+1),2], 1-gcov_cumul[1:maxd],col=col2, type='l', lwd=3,xlab="Depth", ylab="Fraction of regions >= depth",xlim = c(0,20),ylim = c(0,0.01))*

*abline(h = c(0,1,0.5), col = "gray60")*

*abline(v = c(0,50,100,200,300), col = "gray60")*
